# *In vitro* and *in vivo* co-infection and super-infection dynamics of Mayaro and Zika viruses in mosquito and vertebrate backgrounds

**DOI:** 10.1101/2021.06.24.449814

**Authors:** Marco Brustolin, Sujit Pujhari, Cory A. Henderson, Donghun Kim, Jason L. Rasgon

**Affiliations:** Department of Entomology, the Center for infectious Disease Dynamics, and the Huck Institutes of the Life Sciences, the Pennsylvania State University, University Park, PA, USA; The Unit of Entomology, Department of Biomedical Sciences, Institute of Tropical Medicine, Antwerp, Belgium; Department of Pharmacology Physiology and Neuroscience, University of South Carolina School of Medicine, Columbia, South Carolina, USA; Department of Vector Entomology, Kyungpook National University, Daegu, South Korea

## Abstract

Factors related to increasing globalization and climate change have contributed to the simultaneous increase and spread of arboviral diseases. Co-circulation of multiple arboviruses in the same geographic regions provides impetus to study the impacts of multiple arbovirus infections in a single vector. In the present study we describe co-infection and super-infection with Mayaro virus (Family Togaviridae, genus *Alphavirus*) and Zika virus (family Flaviviridae, genus *Flavivirus*) in vertebrate cells, mosquito cells, and *Aedes aegypti* mosquitoes to understand the interaction dynamics of these pathogens and effects on viral infection, dissemination and transmission. In *Aedes aegypti* mosquitoes, co-infection has a negative impact on infection and dissemination rates for Zika virus, but not Mayaro virus, when compared to single infection scenarios, and super-infection of Mayaro virus with a previous Zika virus infection resulted in increased Mayaro virus infection rates. We found that co-infection and super-infection negatively affected Zika viral replication in vertebrate cells (Vero and Huh), resulting in the complete blockage of Zika virus replication in some scenarios. At the cellular level, we demonstrate that single vertebrate and insect cells can be simultaneously infected with Zika and Mayaro viruses. This study highlights the dynamics of arboviral co- and super-infections and emphasizes the importance of considering these dynamics during risk assessment in epidemic areas.

## Introduction

The past several decades have seen dramatic increases in the incidence of arboviral diseases, attributed to increased international trade and transport, climate change, and urban crowding. Increasing landscape fragmentation and anthropic activities are disturbing natural ecosystems, favoring the spill-over of new pathogens with epidemic potential to new geographic regions (1). In the neotropical region of the Americas, more than 150 different arboviruses have been reported (2) and in many regions different arboviruses co-exist and co-circulate in sylvatic and/or urban cycles (3, 4). Despite this diversity and co-circulation of multiple arboviruses in the same geographical area, some pathogens often go unnoticed in both vector and host due to lack of surveillance, overlapping clinical symptoms, and/or the absence of specific diagnostic tests (5). At the same time, clinical studies have identified the co-circulation of multiple arboviruses such as dengue virus (DENV), Zika virus (ZIKV), Chikungunya virus (CHIKV) and Mayaro virus (MAYV) (6–9) in epidemic areas and the presence of multiple arboviruses has been demonstrated in field-collected *Aedes albopictus* (10), supporting the presence and capacity of vectors to carry multiple pathogens simultaneously in nature. Studies under laboratory conditions have also shown that concurrent infection and transmission of multiple arboviruses is possible (11–14).

MAYV (Togaviridae, *Alphavirus*) is an emerging neglected arbovirus thought to be transmitted by *Aedes, Anopheles*, and *Haemagogus* spp. mosquitoes (15). MAYV is actively circulating in South and Central America and was categorized as an emerging threat to public health by PAHO in 2019 (16). ZIKV (Flaviviridae, *Flavivirus*) is primarily transmitted by *Aedes* mosquitoes and emerged as a major arboviral public health concern due to its capacity to cross the placenta, affecting the developing fetus in pregnant women (17, 18). Overlapping distribution and the presence of common vectors (i.e. *Aedes* mosquitoes) in South and Central America (Figure 1) suggest the potential for co-infection among human patients, possibly mediated by the bites of co-infected mosquitoes.

**Figure 1.**
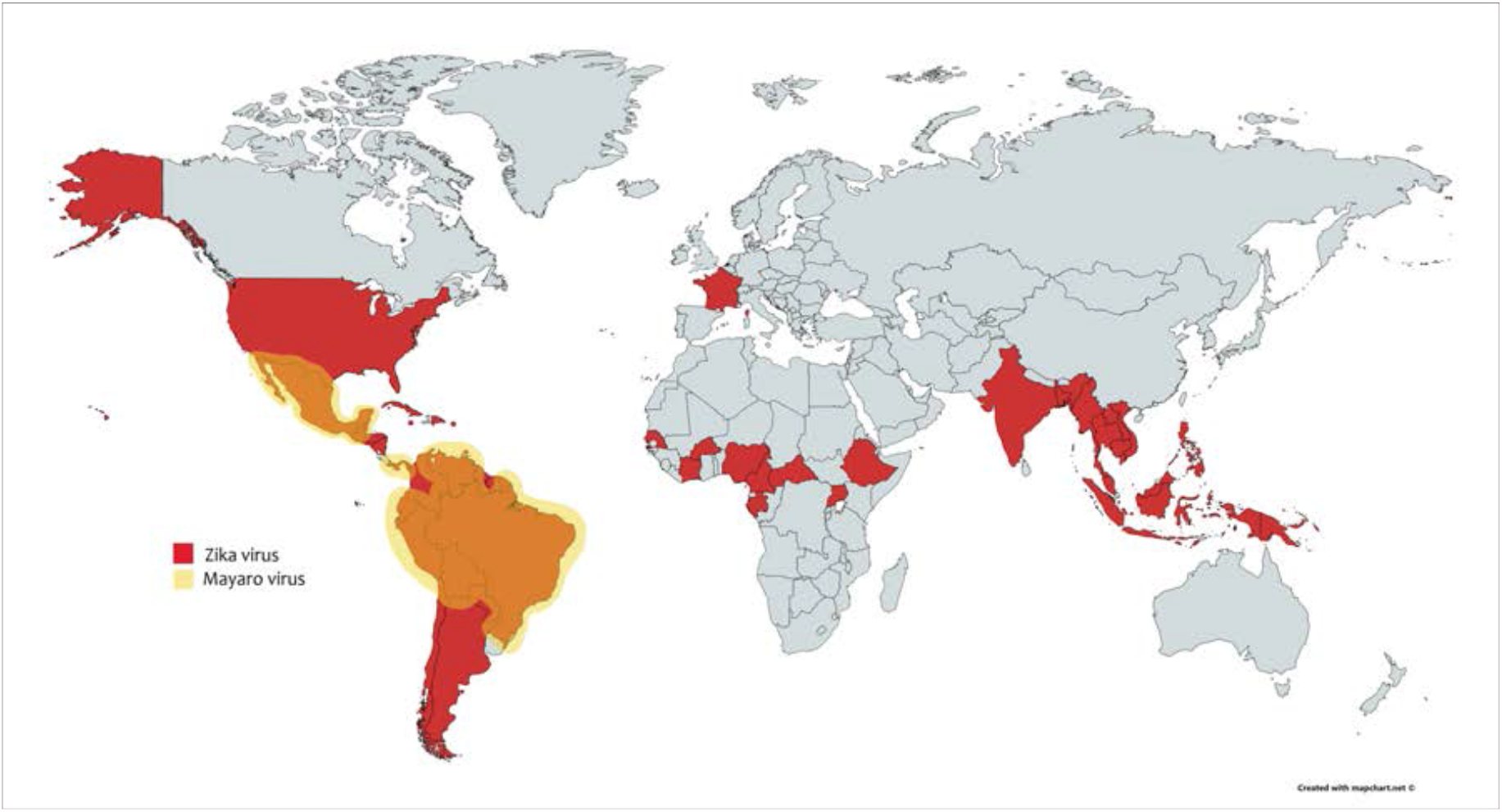
Global distribution of MAYV and ZIKV. Map depicts countries with historical and current detection of MAYV and ZIKV. Map is based on data provided by the CDC as well as (38), and was generated using mapchart.net/detworld.html.

Simultaneous exposure of a vector to multiple pathogens through a single infection event is referred to as co-infection (CI), while serial infection of the vector with different pathogens during subsequent feeding events is referred to as super-infection (SI) (19, 20) (Figure 2). When multiple arboviruses infect the same vector, simultaneously or serially, three different outcomes can occur: 1) an agonistic interaction that results in a partial or total inhibition of one or both viruses, 2) a synergic interaction that enhances one or both viruses or 3) a neutral co-existence of both viruses without any alteration of viral fitness (21). Here, we co-infect and super-infect *Ae. aegypti* mosquitoes with MAYV and ZIKV to study their interactions and the consequent effects on vector competence. We also investigate the interaction kinetics of co-infection and superinfection in vertebrate and invertebrate cells, and we demonstrate that co-infection is possible at the single cell level.

**Figure 2.**
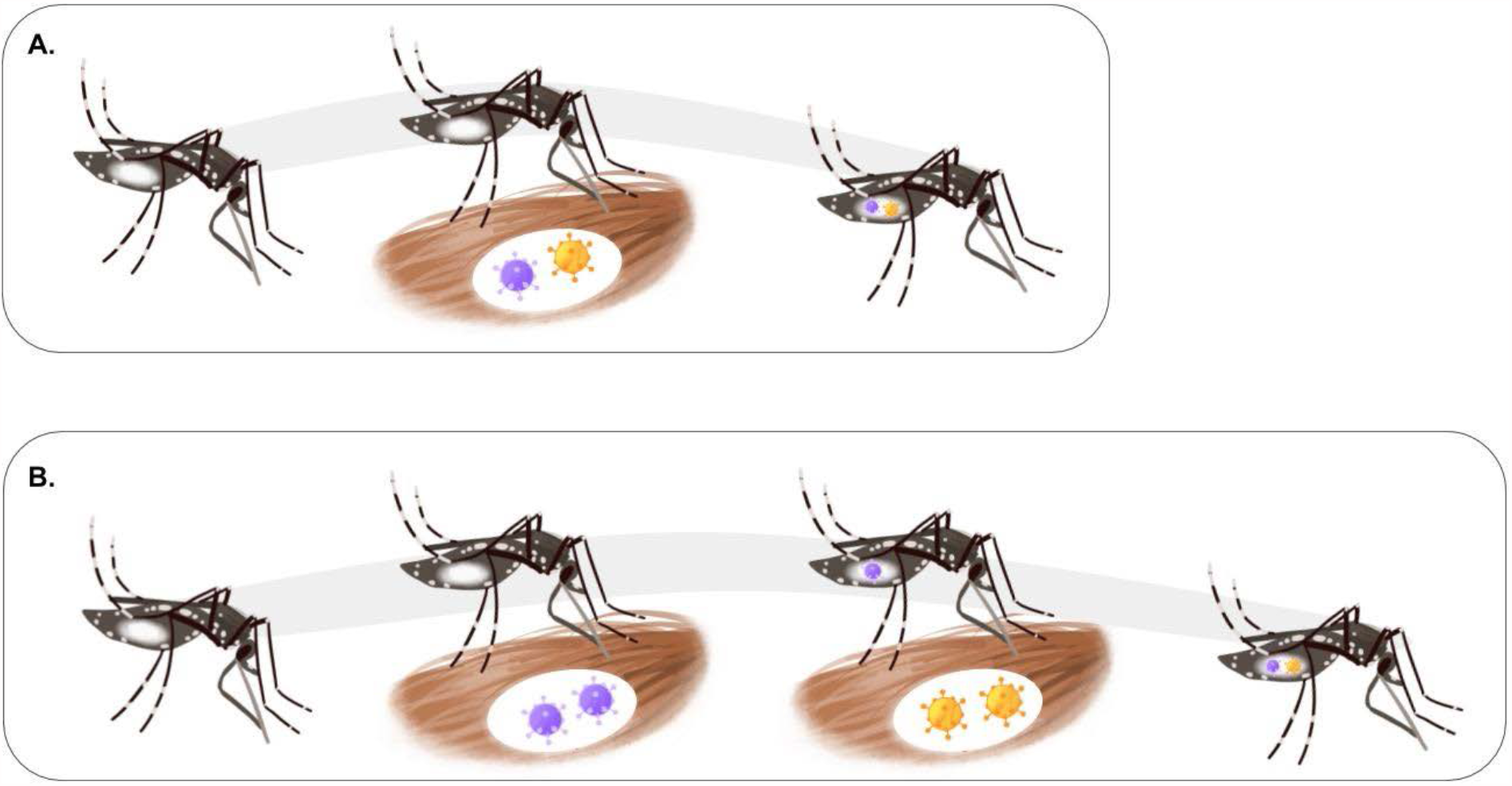
Co-infection and Super-infection in mosquitoes. Competent mosquitoes get infected with multiple arboviruses during a single feeding event (Co-infection; Figure 2a) or sequential feeding events (Super-infection; Figure 2b).

## Results

### Co-infection and Super-infection effects on vector competence and viral titers

*Ae. aegypti* were orally challenged with MAYV and ZIKV simultaneously or serially to investigate the effect of CI and SI on vector competence (Table 1). Infection rate (IR), Dissemination rate (DIR) and Transmission efficiency (TE) were analyzed and compared by quantitative live virus titration in the body, legs, and saliva respectively.

**Table 1.** Infection rate (IR), dissemination rate (DIR), and transmission efficiency (TE) for MAYV and ZIKV in CI and SI experiments. In the co-infection experiment, the IR, DIR and TE have been compared between single infected groups (MAYV [M in the table] or ZIKV [Z in the table]) and the co-infection group (MAYV + ZIKV [CI in the table]). In the SI experiment, the rates of superinfected groups (MZ and ZM) have been compared to control groups fed with different combination of pathogen (M or Z) and Uninfected blood (U) (UM and MU or UZ and ZU). CI= co-infection. M= Mayaro virus. Z= Zika virus. U= Uninfected Blood. P= p-value. ns= not significant. In Superinfection experiment two letters code represent two sequential feeding, ie. MZ= Mayaro virus in the first bloodmeal and Zika virus in the second.

| Coinfection |  |  |  |  |
| --- | --- | --- | --- | --- |
|  | Groups | IR & p-value | DIR & p-value | TE & p-value |
| MAYV positivity |  |  |  |  |
| 7dpi | M vs CI | 14/23 (60%) vs 9/19 (47%); ns | 13/14 (92%) vs 7/9 (78%); ns | 1/23 (4%) vs 0/19; ns |
| 14dpi | M vs CI | 17/23 (73%) vs 12/19 (63%); ns | 17/17 (100%) vs 11/12 (92%); ns | 1/22 (4%) vs 2/19; ns |
| ZIKV positivity |  |  |  |  |
| 7dpi | Z vs CI | 25/25 (100%) vs 14/19 (73%); p= 0.0107 | 18/25 (72%) vs 4/14 (28%); p= 0.0019 | 0/25 vs 0/19; ns |
| 14dpi | Z vs CI | 25/26 (96%) vs 12/19 (63%); p= 0.0064 | 24/25 (96%) vs 11/12 (92%); ns | 8/26 (31%) vs 3/19 (16%); ns |
| Superinfection |  |  |  |  |
| MAYV positivity |  |  |  |  |
| 7dpi | MU vs MZ | 12/12 (100%) vs 4/5 (80%); ns | 12/12 (100%) vs 3/4 (75%); ns | 3/12 (25%) vs 0/5; ns |
| 14dpi | MU vs MZ | 12/12 (100%) vs 7/8 (87%); ns | 10/12 (83%) vs 6/7 (85%); ns | 1/12 (8%) vs 1/8 (12%); ns |
| 7dpi | UM vs ZM | 9/14 (64%) vs 20/21 (95%) p= 0.0278 | 8/9 (88%) vs 18/20 (90%); ns | 2/14 (14%) vs 0/21; ns |
| 14dpi | UM vs ZM | 13/14 (93%) vs 18/22 (81%); ns | 13/13 (100%) vs 17/18 (94%); ns | 0/12 vs 1/22 (4%); ns |
| ZIKV positivity |  |  |  |  |
| 7dpi | ZU vs ZM | 14/14 (100%) vs 21/21 (100%); ns | 14/14 (100%) vs 21/21 (100%); ns | 9/14 (64%) vs 10/21 (48%); ns |
| 14dpi | ZU vs ZM | 11/11 (100%) vs 23/23 (100%); ns | 10/11 (90%) vs 20/23 (87%); ns | 7/11 (63%) vs 8/23 (35%); ns |
| 7dpi | UZ vs MZ | 20/20 (100%) vs 5/0 (100%); ns | 17/20 (85%) vs 4/5 (75%); ns | 0/20 vs 0/5; ns |
| 14dpi | UZ vs MZ | 17/17 (100%) vs 8/8 (100%); ns | 17/17 (100%) vs 8/8 (100%); ns | 9/17 (53%) vs 5/8 (62%); ns |

In the CI experiment, we observed a significant reduction in IR for ZIKV in the co-infected group compared to the single infected group at 7 and 14 days post-infection (dpi) (100% vs 73% p= 0.0107 and 96% vs 63% p= 0.0064 respectively) (Table 1). Similar reduction was also observed for the DIR at 7 dpi (72% vs 28% p=0.0019) (Table 1). We did not observe any statistically significant difference in IR or DIR for MAYV in the co-infection group in comparison to the single infected group at any time point (Table 1).

Zika virus titer in the body and legs of co-infected mosquitoes was statistically lower compared to the titer recorded in the body and legs of single infected mosquitoes at 7 dpi (body p< 0.0001; legs p< 0.0001) and 14 dpi (body p= 0.0002; legs p= 0.0041) (Figure 3 panel A graphs 1-4). Similarly, MAYV titers showed a significant reduction in the body of co-infected mosquitoes at 14 dpi (p=0.0130) (Figure 3 panel A graph 2).

**Figure 3.**
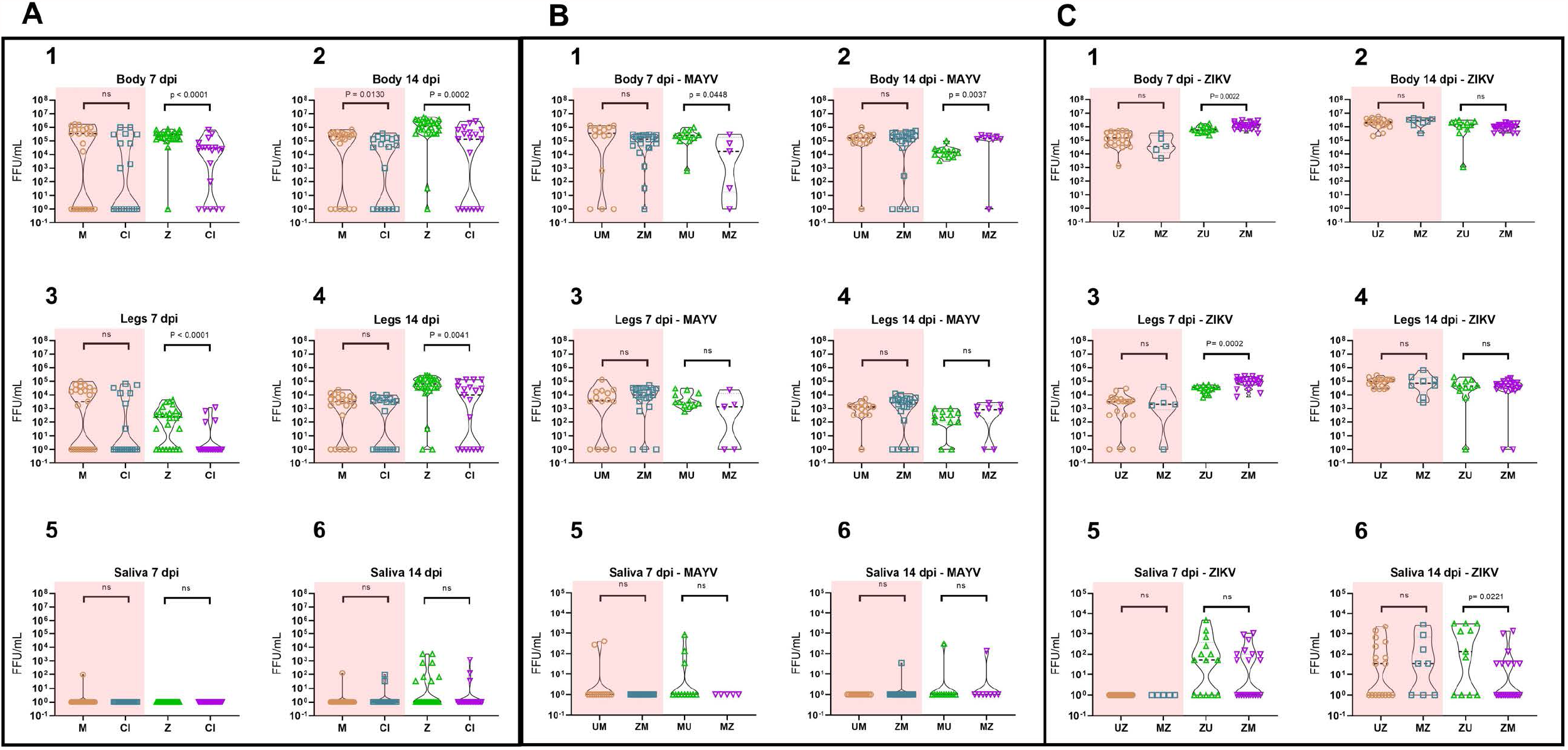
Infectious virus loads of MAYV and ZIKV in co-infection and super-infection Ae. aegypti. Panel A shows MAYV and ZIKV viral loads in the co-infection experiment. Titers obtained from single infected groups were compared with those from the co-infected group. Panel B shows the viral loads of MAYV in the super-infection experiment. In this panel, two different comparisons were made: UM vs ZM and MU vs MZ. Panel C shows the viral loads of ZIKV in the super-infection experiment. In this panel two different comparisons were done: UZ vs MZ and ZU vs ZM. Titers are expressed in FFU/mL. CI= co-infection. M= Mayaro virus. Z= Zika virus. U= Uninfected Blood. p= p value. ns= not significant.

We did not find any statistically significant differences in the TEs or respective saliva viral titers of MAYV or ZIKV between single infected and co-infected groups. These data suggest that the simultaneous intake of both viruses has a negative impact on their respective capacity to infect the midgut and the subsequent spread to the whole body of the mosquito, but not on the overall capacity of the mosquitoes to transmit both viruses in accordance to what is previously demonstrated for CHIKV/ZIKV co-infection (12, 13).

The SI experiment is designed to mimic the most common scenario in nature for mosquito species that blood feed multiple times during a single gonotrophic cycle, as the case of *Ae. aegypti* or *Ae. albopictus*. In this experiment we recorded only one statistically significant difference for IR, DIR or TE among all the combinations tested: the increase of the IR for MAYV in the ZM group compared to the UM group (95% vs 64% p= 0.0278) at 7 dpi. This result indicates a positive effect of a previous ZIKV infection on MAYV ability to establish a stable infection in the midgut.

The enhancement of MAYV IR caused by ZIKV did not lead to higher MAYV titers in the body. We did not observe any statistically significant variation of MAYV titer in mosquitoes previously exposed to ZIKV compared to those previously exposed to uninfected blood. Conversely, our results indicate that subsequent ZIKV infection can influence MAYV titer. We observed a statistically significant decrease of MAYV titer in body of the MZ group at 7 dpi (group MZ; p= 0.0448; Figure 3 panel B graph 1) however, the same experimental group showed an opposite behavior at 14 dpi, when a statistically significant increase of MAYV body titer was detected (group ZM p= 0.0037; Figure 3 panel B graph 2).

The titer of ZIKV was positively affected by a subsequent MAYV infection at 7 dpi (Figure 3 panel C graphs 1 and 3). We observed a statistically significant enhancement of ZIKV titer in the body and legs of the ZM group compared to the ZU group (p=0.0022 and p=0.0002 respectively) at 7 dpi. These differences disappear at 14 dpi where the titers are similar in both body and legs.

Saliva titers of ZIKV showed a reduction in viral titer in the saliva of the ZM group compared to the ZU group at 14 dpi (p = 0.0221; Figure 3 panel C graph 6).

In the CI experiment we observed similar IR for double positive mosquitoes at 7 and 14 dpi (7/19 (37%)) (Table 2). Conversely, mosquitoes with double disseminated infection were observed only at 14 dpi (6/19 (32%)). No transmission was observed at any time point for the CI group.

**Table 2.** Infection rate (IR), dissemination rate (DIR), and transmission efficiency (TE) for double positive mosquitoes in CI and SI experiments. In the SI experiment, the rates of superinfected groups (MZ and ZM) have been compared with Fisher’s exact test at different time points. CI= co-infection. M= Mayaro virus. Z= Zika virus. U= Uninfected Blood. * = DIR was calculated as number of mosquitoes with double positive legs/ total number of exposed mosquitoes. In Superinfection experiment two letter codes represent two sequential feeding, (ie. MZ= Mayaro virus in the first bloodmeal and Zika virus in the second).

|  |  | Coinfection |  |  |
| --- | --- | --- | --- | --- |
|  | Groups | IR – Double positive | DIR - Double positive | TE - Double positive |
| 7dpi | CI | 7/19 (37%) | 0/19* | 0/19 |
| 14dpi | CI | 7/19 (37%) | 6/19* (32%) | 0/19 |
|  |  | Superinfection |  |  |
| 7dpi | MZ | 4/5 (80%) | 3/5* (60%) | 0/5 |
| 14dpi | MZ | 7/8 (86%) | 6/8* (75%) | 1/8 (12%) |
| 7dpi | ZM | 20/21 (95%) | 18/21* (86%) | 0/21 |
| 14dpi | ZM | 18/22 (82%) | 16/22* (73%) | 1/22 (5%) |

In the SI experiment, we observed higher IR of double positive mosquitoes compared to CI experiment (Table 2). In addition, double disseminated infection was recorded starting from 7 dpi, contrary to what was observed in the CI experiment. These data suggest that vectors are more permissive to double infection with multiple arboviruses in case of subsequent exposure rather than from simultaneous exposure. Saliva samples tested positive for both viruses only at 14 dpi. In the ZM group, 1/22 (4.5%) saliva samples tested positive for both viruses; in the MZ group 1/8 (12.5%) saliva samples tested positive for both viruses. Comparisons between IR, DIR and TE of MZ and ZM groups did not show any statistically significant difference.

Finally, we analyzed and compared titers of double positive body and legs, but no statistically significant results between MAYV and ZIKV titers were found in both CI and SI experiments.

### Growth kinetics of MAYV and ZIKV during single, co-infection and super-infection

To determine if CI or SI results in viral growth inhibition, we performed growth-curve analysis of MAYV and ZIKV in Aag2 cells (derived from *Ae. aegypti*; mosquito vector), Vero cells (derived from African green monkey; primate reservoir), and Huh7.5 cells (derived from human; human host) infected with one or both viruses at 0.1 MOI. MAYV and ZIKV infectious virus production were quantified in the culture supernatants (Figure 4).

**Figure 4.**
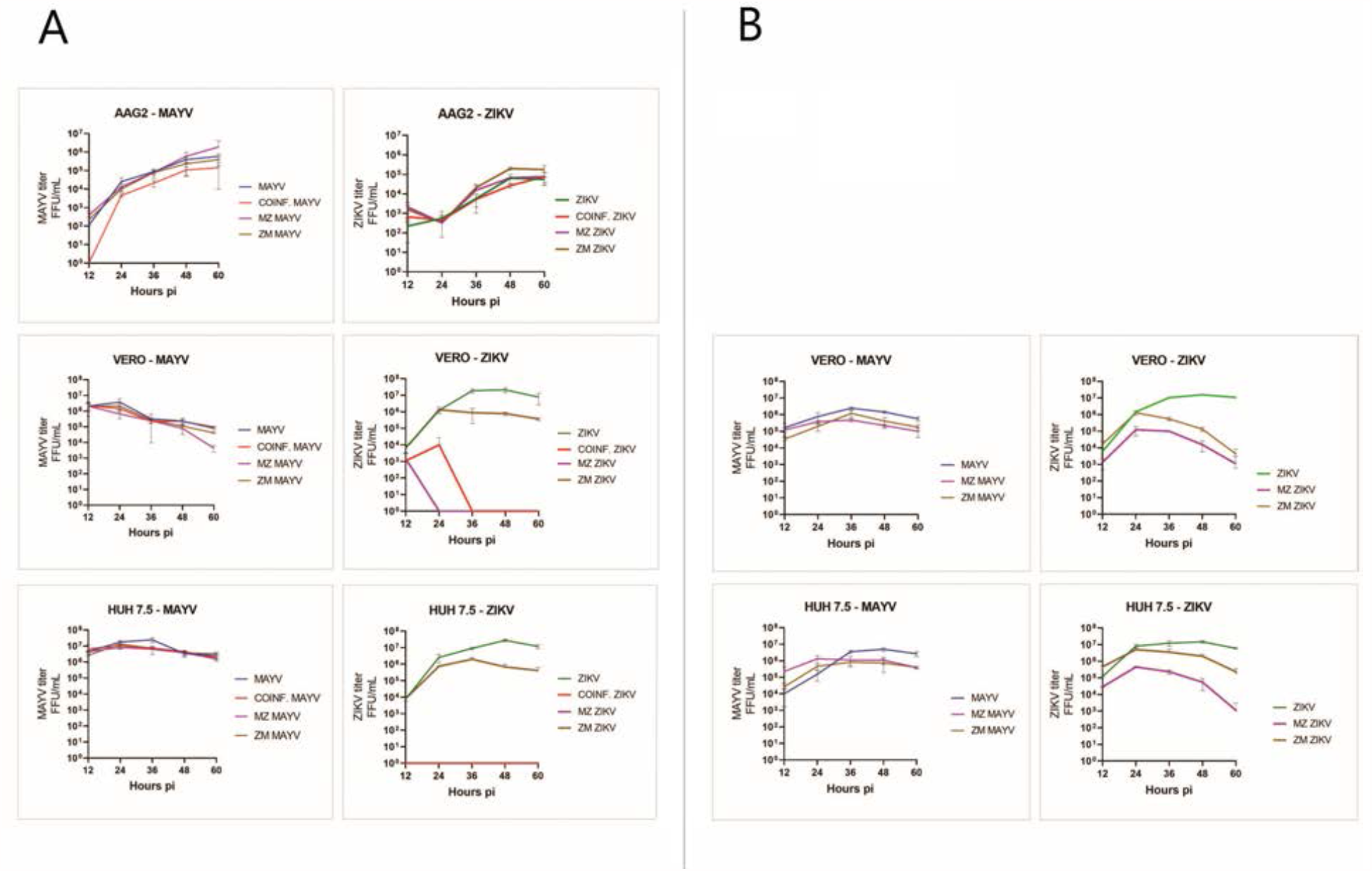
Mayaro and Zika virus growth curves in select cell types at different time points and experimental infection conditions. Aag2, Huh7.5 and Vero cells were infected with MAYV and ZIKV at 0.1 MOI as mono-, co- or sequential infection. A) cells were superinfected after 12-hours post infection; B) vertebrate cells were superinfected after 2-hours post infection. Culture supernatants were collected every 12h and virus was titrated and shown as FFU/ml. Data presented from three technical replicates. Log10-transformed viral titers were analyzed using R and GraphPad.

In single-infected Aag2 cells, MAYV titer continued to increase after 60h post-infection (5.5 log/ml) while ZIKV titer reached the peak at 48h (4.5 log/ml) followed by a plateau phase. In vertebrate cells, viral titers reached a peak before decreasing. MAYV titers reached their peak 24h post-infection in Vero (6.3 log/ml) and 36h post-infection in Huh7.5 cells (7.2 log/ml); ZIKV titers reached their peak 48h post-infection both in Vero and Huh7.5 cells (7.2 log/ml). The peak of MAYV titer was one log lower in Vero cells and two logs lower in Aag2 cells compared to the Huh7.5 cells. ZIKV kept producing infectous virus until 60h post-infection in Aag2 cells, while in Vero and Huh7.5 cells it reached peak titer at 48h post-infection with a titer of 7.2 log/ml (Figure 4).

Co-infection and super-infection had significant effects on the replication and release of infectious virus particles, primarily ZIKV in vertebrate cells. In both CI and SI post MAYV infection, ZIKV replication and production of infectious virus particles were completely blocked in Vero and Huh7.5 cells, indicating a significant competition or inhibition of ZIKV replication by MAYV (Figure 4A). In order to investigate this phenomenon, we repeated the SI experiment in vertebrate cells adding the second virus 2 hours after the first as opposed to 12 hours. Interestingly, when infected with ZIKV at 2-hour post infection of MAYV, vertebrate cells could support replication and production of Zika virus at lower levels (Figure 4B).

### MAYV and ZIKV can simultaneously infect and replicate in the same cell

To determine the capacity of MAYV and ZIKV to infect and replicate in the same cell simultaneously, we immunolocalized both viruses in Aag2 and Vero cells (Figure 5) using immunofluorescence microscopy. MAYV, like other alphaviruses, was predominantly localized to the cell membrane and filapodial extensions. Due to massive and rapid cytopathic effect caused by MAYV, most infected vertebrate cells showed the formation of membrane protrusions. Similar cellular structures and virus distribution were identified previously in CHIKV infected cells (13). ZIKV was primarily localized on the perinucleus and endoplasmic reticulum regions, the proposed replication and assembly sites for flaviviruses (22). A similar pattern of distribution of MAYV and ZIKV in Aag2 cells indicates both these viruses can co-infect the same cell in their vertebrate and invertebrate host.

**Figure 5.**
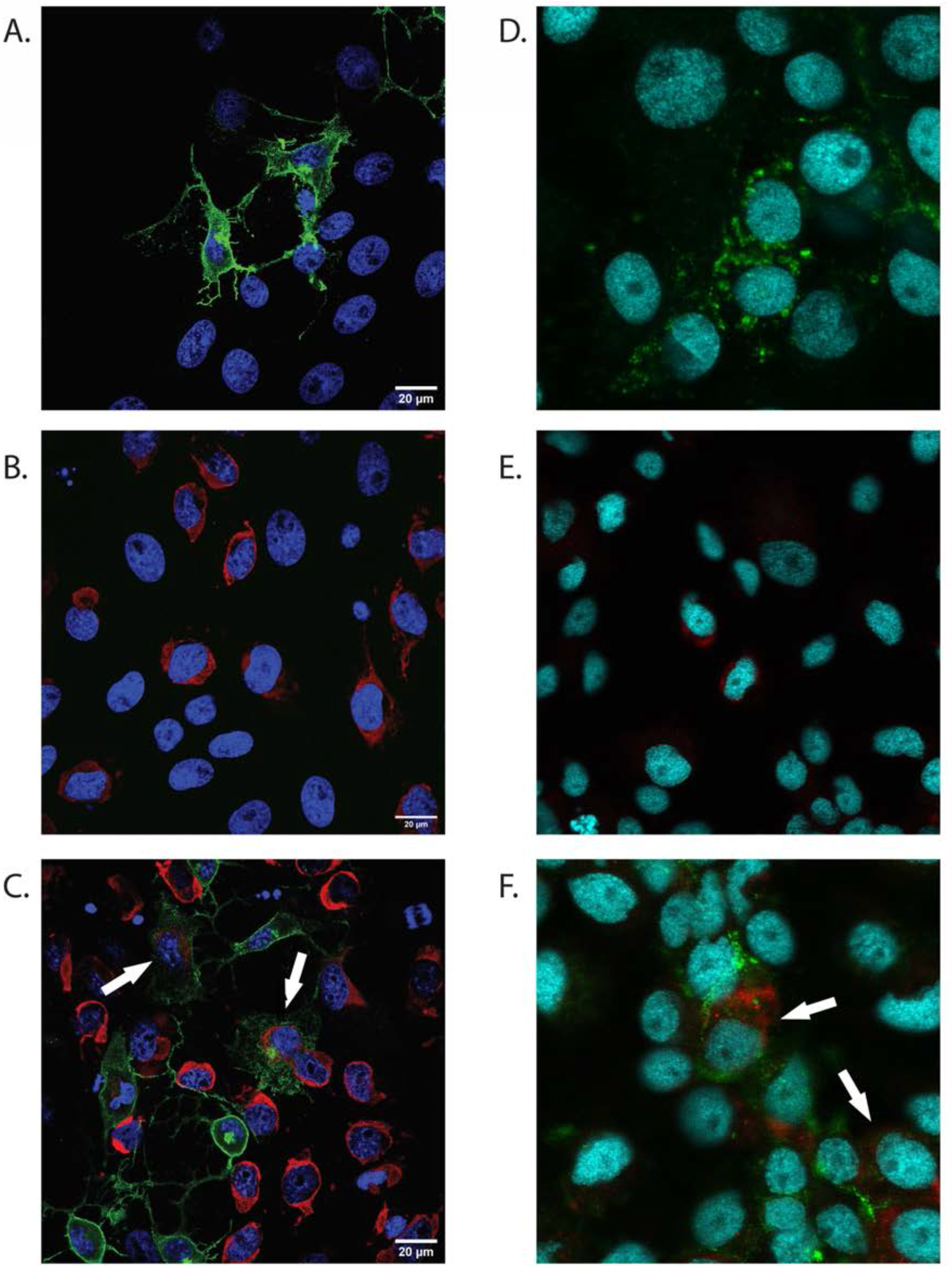
Mayaro and Zika virus can infect and replicate in the same cell. (Co-) Infection of Aag2 and Vero cells was performed at 1 MOI. Infected monolayers were fixed after 24h (Vero cells) or 48h (Aag2 cells) and labeled with monoclonal anti-CHIKV E2 envelope glycoprotein clone CHK-48 (MAYV) or/and polyclonal anti-Zika virus envelope protein MAYV = green. ZIKV = red. Cell Nuclei = Blue. White arrows indicate co-infected cells.

## Discussion

Here we show that *Ae. aegypti* can be infected with MAYV and ZIKV simultaneously after sequential or co-infection with both pathogens and that simultaneous transmission of MAYV and ZIKV by *Ae. aegypti* is possible as a result of a super-infection event. Even though there are no data on MAYV/ZIKV co-infected mosquitoes in nature, pools of mosquitoes collected in different areas of Mato Grosso (Brazil) were double-positive for MAYV and DENV-4 (23). In addition, reports of MAYV co-infected patients with CHIKV (8), ZIKV (24), and DENV-1 (25), indicate that MAYV is co-circulating in urban and peri-urban areas, probably due to the high number of vector species that can contribute to its transmission cycle (15) resulting in potentially double infected mosquitoes. Currently, most surveillance programs pool and analyze mosquitoes to save time and resources. Unfortunately, this surveillance strategy makes it impossible to investigate the percentage of the mosquitoes carrying multiple arboviruses and their potential contribution to the transmission cycle of those viruses. The implementation of precise sampling strategies of field-collected mosquitoes would be fundamental to investigate the prevalence of multi-infected vectors in the endemic areas and correlate the data with the incidence of co-infected patients.

The dynamics of co-infection and super-infection of MAYV and ZIKV at cellular and molecular levels are still unclear. The results obtained *in-vitro* demonstrate that dual infection of these viruses both in invertebrate and vertebrate cells are possible. However, we noticed several cases of viral interference. MAYV inhibits ZIKV replication in mammalian cells in the case of CI and of previous infection (MZ). At the same time, only partial inhibition was recorded in sequential infection (ZM). Interesting, when the SI experiment in mammalian cells was repeated with an interval of 2h (instead of 12h) incubation period between sequential infections, ZIKV could replicate in MAYV infected cells. The inability of entry or replication of the second virus in the presence of another virus is also called viral interference. Interference of arbovirus replication mediated by another arbovirus is influenced by several principal parameters: advantage time of the first virus time, the exposure order of viral pathogens, and difference in MOI (26). Different hypothesis haves been proposed to explain viral interference mechanisms including competition for replication sites and cellular substrate resources (26), trans-acting proteases induced by the first virus, and super-infection exclusion (27, 28). Super-infection exclusion (SIE) occurs when a cell infected by a virus becomes refractory to subsequent infection by the same or a closely related virus (27) and can occurs at different step of replication cycle, including entry, viral genome transcription and protein translation (29). In a recent study, Boussier et al. showed that CHIKV excludes influenza A virus (Orthomyxoviridae, *Alphainfluenzavirus*) replication *in vitro* and demonstrated the capacity of alphaviruses to interfere with replication cycle of different viral families (29). Several *in vivo* studies suggest that sequential infection of *Alphavirus* and *Flavivirus* results in co-infection, not in exclusion (13, 30, 31), however none of them used MAYV. Here we showed the presence of co-infected cells using immunofluorescence microscopy, proving that a single cell can be simultaneously infected with both viruses (Figure 5). We observed a cytopathic effect in mammalian cells caused by MAYV after the first 12-24 hours post infection, similar to what previously described for other alphaviruses (27). The loss of structural functionality and the partial destruction of the cells might have partially prevented the cell’s infection by ZIKV and its consequent replication, especially in case of long incubation time between infections (12h). The replication cycle of alphaviruses is relatively short and progeny virions can be detected 4-6 hours post infection (32–34). Conversely, progeny virions of flaviviruses can be generally detected 8-12 hours post infection (35). This fitness disparity could have influenced the capacity of ZIKV to replicate in presence of MAYV, which rapidly induce a severe structural and morphological change in the infected cells. Super-infection with short incubation time (2h) has probably limited the impact of cell damage caused by the first infection, however in all the experimental conditions MAYV inhibits ZIKV replication to some degree. We can therefore speculate that when MAYV is introduced in vertebrate cells before or together with ZIKV, it takes over the cellular transcription and/or translational machinery leaving minimal resources for replication of ZIKV. Additional studies are required to understand the molecular dynamics of CI and SI of MAYV and ZIKV in both invertebrate and vertebrate models.

Our results *in vivo* showed different outcomes for IR and DIR of MAYV and ZIKV in case of co-infection. We observed a significant reduction in the IR of ZIKV during a co-infection. Rückert et *al*. demonstrated a reduction of IR for ZIKV in the case of CI with CHIKV (12), an *Alphavirus* belonging to the same antigenic complex of MAYV (Semliki Forest virus complex). Similarly, Muturi et *al*. demonstrated that in co-infected *Ae. albopictus*, replication of DENV-4 was suppressed (20). We did not observe any variation in the IR of MAYV during co-infection with ZIKV. Our data are consistent with previous studies using different a *Flavivirus/Alphavirus* co-infection models. In these studies, the authors demonstrated that IR and DIR of CHIKV were not influenced by the presence of ZIKV or DENV-2 (12, 36). Taken together, these data suggest that in co-infected mosquitoes the IR of flaviviruses is negatively affected by the presence of an *Alphavirus* but not the opposite.

Interestingly, we observed an increase of MAYV IR in previously ZIKV-infected mosquitoes (ZM group). The molecular mechanism underlying this finding is not clear; however, we speculate that the previous ZIKV infection could have altered the mosquito immune response, creating a more suitable environment for MAYV, similar to what has been hypothesized for CHIKV-ZIKV super-infection (14). Our experimental model assumes that the vector gets co-infected or super-infected with a similar titer of both viruses - a scenario that may not reflect what happens in the field where the possible combination of viruses and their respective titers are incredibly variable. Muturi *et. al*. (20) demonstrated that varying virus titer ratio between DENV-4 and SINV could affect the replication of one or both viruses in an *in vitro* SI study, suggesting that the relative amounts of different pathogens could influence the outcome of CI or SI experiments. Therefore, additional studies are required to evaluate the impact of different titer combinations in case of CI or SI.

## Material and Methods

### Mosquito

*Ae. aegypti* (Rockefeller strain) was obtained from Johns Hopkins University. Mosquito colonies were reared and maintained at the Millennium Sciences Complex insectary (The Pennsylvania State University, University Park, PA, USA) under the following environmental conditions: 27°C ± 1°C, 12:12 h light:dark diurnal cycle at 80% relative humidity. The larvae were fed with ground fish flakes (TetraMin, Melle, Germany). Adult mosquitoes had free access to 10% sucrose solution.

### Cells

Vero African green monkey kidney cells (CCL-81; ATCC, Manassas, VA, USA) and Huh7.5 (human liver cell line) (kind gift from Dr. Craig Cameron) were cultured in DMEM (Life Technologies) containing 10% Fetal Bovine Serum (FBS), penicillin (100 U/ml), streptomycin (100 μg/ml), 10 mM HEPES, and 200 mM glutamine. Aag2 *Ae. aegypti* cells (kind gift from Dr. Elizabeth McGraw) were grown in Schneider’s insect medium supplemented with 10% FBS, penicillin (100U/ml), streptomycin (100 μg/ml), and 200 mM glutamine.

### Virus

Zika virus strain MR766 (NR-50065; BEI Resources, Manassas, VA, USA) and Mayaro virus strain BE AR 505411 (NR-49910; BEI Resources, Manassas, VA, USA) were propagated on Vero cells, stocks solutions were aliquoted and stored at −70°C until used. Viral stock titers were obtained by focus forming assay (FFA).

#### *In-vitro* mono-, Co-, and Super-infection

Cells were infected with ZIKV and/or MAYV with a multiplicity of infection (MOI) of 0.1. For mono-infection, cells were incubated independently with either ZIKV or MAYV, together for co-infection or sequentially for super-infection. After 1h of incubation at 37°C, the virus inoculum was removed, cells washed twice to remove unbound virus, and fresh medium added. At each time point, cell-free culture supernatants were collected and stored at −80°C and subsequently analyzed for virus titer by focus forming assay as previously described (15). Briefly, 30 µl of 10-fold serial dilutions of each sample were used to infect a Vero cell monolayer in a 96 well plate. After 24 hours (for MAYV) or 48 hours (for ZIKV), cells were fixed, permeabilized and labeled using the monoclonal anti-CHIKV E2 envelope glycoprotein clone CHK-48 (which reacts with MAYV) (BEI Resources, Manassas, VA, USA) or the monoclonal anti-Flavivirus group antigen (Clone D1-4G2-4-15). CI and SI samples were analyzed twice, one for each specific antibody. Subsequently, the primary antibody was labeled with the Alexa-488 goat anti-mouse IgG secondary antibody (Invitrogen, Life Science, Eugene OR, USA) and fluorescent foci observed and counted with an Olympus BX41 microscope equipped with an UPlanFI 4× objective and a FITC filter. SI experiment in vertebrate cells were performed twice, using two different incubation times between the first and the second infection: 2- or 12-hours. Three tecnical replicates were performed for each condition.

#### Vector competence assay

Unfed 5-7 days old female mosquitoes were separated in groups of approximately 80 individuals and fed with virus-spiked infected human blood via a glass feeder jacketed with 37°C distilled water for 30-45 min (final titer for each virus was 1*10^7 FFU/mL). In the CI experiment blood containing MAYV, ZIKV, or both viruses were presented to different mosquito groups (henceforth called M, Z and CI) (day 0). A control group fed with uninfected blood (U) was included as the negative control. After feeding, mosquitoes were anesthetized at 4°C, engorged females were selected and transferred to a clean cardboard cup with access to 10% sugar solution *ad libitum*. In the SI experiment, two sequent feeding events were scheduled. During the first feeding operation (day-6) 3 groups were exposed to uninfected blood, 2 groups to ZIKV (Z) and 2 groups to MAYV (M). During the second feeding operation (day 0) all the groups were exposed to a different bloodmeals resulting in 7 different possible combinations: UM, UZ, MU, ZU, MZ, ZM and UU. After both feedings, full-engorged females were selected and housed has previously described.

At 7 and 14 days post-infection (dpi), living mosquitoes were anesthetized with triethylamine (Sigma, St. Louis, MO) for approximately 30 seconds and subsequently dissected by separating the legs from the body of the mosquito. After leg detachment, the mosquitoes were forced to salivate in a glass capillary filled with a 1:1 solution of 50% sucrose solution and FBS for 30 minutes. Bodies and legs were collected in separate 2mL tube containing 1 ml of mosquito diluent (20% FBS in Dulbecco’s phosphate-buffered saline, 50 µg/ml penicillin/streptomycin, 50 ug/ml gentamicin, and 2.5 µg/ml fungizone) with a single sterile zinc-plated, steel 4.5 mm BB (Daisy, Rogers, AR, USA). Tissues were homogenized at 30 Hz for 2 min using TissueLyser II (Qiagen GmbH, Hilden, Germany) and centrifuged for 30 sec at 11000 rpm. Saliva samples were collected in a 2ml tube containing 0.1 ml of mosquito diluent. All samples were stored at −70°C until used. Samples were analyzed by Focus Forming assay as described above.

The experimental infections were performed at the Eva J. Pell ABSL-3 Laboratory for Advanced Biological Research (Pennsylvania State University) Infection rates (IR) (positive bodies/total of exposed mosquitoes), dissemination rates (DIR) (positive legs/positives bodies), and transmission efficiency (TE) (positive saliva/ total of exposed mosquitoes) were calculated for all the tested groups. In the co-infection experiment we compared M vs CI and Z vs CI. In the super-infection experiment we compared MU vs MZ, ZU vs ZM, UM vs ZM, and UZ vs MZ. The titers of MAYV and ZIKV in body, legs and saliva of each mosquito in each group were calculated and expressed in FFU/mL. Viral titers in specific tissues were compared between group following the scheme used for the comparison of IR, DIR and TE and described in the previous paragraph.

#### Immunolocalization of MAYV and ZIKV on Vero and Aag2 cells

Cells were seeded at a density of 1 × 10^5^ cells/well (Vero cells) or 1 × 10^6^ cells/well (Aag2 cells) in a 2-well chamber slide and grown at 37°C with 5% CO_2_ (Vero cells) or 27 °C without CO_2_ (Aag2). After 24 hours, cell monolayers were infected with ZIKV, MAYV or both at a MOI of 1. After 24 (Vero cells) or 48 (Aag2) hours, cells were fixed and permeabilized following the same procedure described above. Then cells were labeled with the monoclonal anti-CHIKV E2 envelope glycoprotein clone CHK-48 made in mouse (BEI Resources, Manassas, VA, USA) and the polyclonal anti-Zika virus envelope protein made in rabbit (GeneTex, Irvine, CA, USA). The antibodies used for immunofluorescence included Alexa Fluor 488 conjugated goat anti-rabbit IgG (Invitrogen) and Alexa Fluor 594 conjugated donkey anti-rabbit IgG (Invitrogen). Hoechst 33342 Fluorescent Stain (Thermo Scientific) was used for nuclear staining. Vero cells were examined using an Olympus ZEISS LSM 900 confocal microscope (Zeiss, Germany), Aag2 cells using a Olympus Fluoview 10i-LIV confocal microscope (Olympus Corporation, US), and images processed using ImageJ software (37).

### Statistical analysis

GraphPad Prism software version 8.2.1 (441) was used to analyze all the data. Differences in the IR, DIR and TE of groups challenged in both experiments were analyzed by Fisher’s exact test. Two-tailed Mann-Whitney U test was used to compare viral titers in body, legs, and saliva samples of different groups. A p value of < 0.05 was considered statistically significant.

## Acknowledgments

We thank the personnel of Eva J. Pell ABSL-3 Laboratory for Advanced Biological Research for their help and technical support, Drs. Craig Cameron and Elizabeth McGraw for cell lines, and Sage McKeand for Figure 2.

